# A high-resolution, chromosome-assigned Komodo dragon genome reveals adaptations in the cardiovascular, muscular, and chemosensory systems of monitor lizards

**DOI:** 10.1101/551978

**Authors:** Abigail L. Lind, Yvonne Y.Y. Lai, Yulia Mostovoy, Alisha K. Holloway, Alessio Iannucci, Angel C.Y. Mak, Marco Fondi, Valerio Orlandini, Walter L. Eckalbar, Massimo Milan, Michail Rovatsos, Ilya G. Kichigin, Alex I. Makunin, Martina J. Pokorná, Marie Altmanová, Vladimir A. Trifonov, Elio Schijlen, Lukáš Kratochvíl, Renato Fani, Tim S. Jessop, Tomaso Patarnello, James W. Hicks, Oliver A. Ryder, Joseph R. Mendelson, Claudio Ciofi, Pui-Yan Kwok, Katherine S. Pollard, Benoit G. Bruneau

## Abstract

Monitor lizards are unique among ectothermic reptiles in that they have a high aerobic capacity and distinctive cardiovascular physiology which resembles that of endothermic mammals. We have sequenced the genome of the Komodo dragon (*Varanus komodoensis*), the largest extant monitor lizard, and present a high resolution *de novo* chromosome-assigned genome assembly for *V. komodoensis*, generated with a hybrid approach of long-range sequencing and single molecule physical mapping. Comparing the genome of *V. komodoensis* with those of related species showed evidence of positive selection in pathways related to muscle energy metabolism, cardiovascular homeostasis, and thrombosis. We also found species-specific expansions of a chemoreceptor gene family related to pheromone and kairomone sensing in *V. komodoensis* and several other lizard lineages. Together, these evolutionary signatures of adaptation reveal genetic underpinnings of the unique Komodo sensory, cardiovascular, and muscular systems, and suggest that selective pressure altered thrombosis genes to help Komodo dragons evade the anticoagulant effects of their own saliva. As the only sequenced monitor lizard genome, the Komodo dragon genome is an important resource for understanding the biology of this lineage and of reptiles worldwide.

## Introduction

The evolution of form and function in the animal kingdom contains numerous examples of innovation and diversity. Within vertebrates, non-avian reptiles are a particularly interesting lineage. There are an estimated 10,000 reptile species worldwide, found on every continent except Antarctica, with a diverse range of morphologies and lifestyles ^1^. This taxonomic diversity corresponds to a broad range of anatomic and physiological adaptations. Understanding how these adaptations evolved through changes to biochemical and cellular processes will reveal fundamental insights in areas ranging from anatomy and metabolism to behavior and ecology.

The varanid lizards (genus *Varanus*, or monitor lizards) are an unusual group of squamate reptiles characterized by a variety of traits not commonly observed within non-avian reptiles. Varanid lizards vary in mass by close to five orders of magnitude (8 grams–100 kilograms), comprising the genus with the largest range in size ^2^. Among the squamate reptiles, varanids have a unique cardiopulmonary physiology and metabolism with numerous parallels to the mammalian cardiovascular system. For example, their cardiac anatomy allows high pressure shunting of oxygenated blood to systemic circulation ^3^. Furthermore, varanid lizards can achieve and sustain very high aerobic metabolic rates accompanied by elevated blood pressure and high exercise endurance ^4–6^, which facilitates high-intensity movements while hunting prey ^7^. The specialized anatomical, physiological, and behavioral attributes of varanid lizards are all present in the Komodo dragon (*Varanus komodoensis*). As the largest extant lizard species, Komodo dragons can grow to 3 meters in length and run at speeds of up to 20 kilometers miles per hour, which allows them to hunt large prey such as deer and boar in their native Indonesia ^8^. Komodo dragons have a higher metabolism than predicted by allometric scaling relationships for varanid lizards ^9^, which helps explains their extraordinary capacity for daily movement to locate prey ^10^. Their ability to locate injured or dead prey through scent tracking over several kilometers is enabled by a powerful olfactory system ^8^. Additionally, serrate teeth, sharp claws, and saliva with anticoagulant and shock inducing properties aid Komodo dragons in hunting prey ^11,12^. Komodo dragons have aggressive intraspecific conflicts over mating, territory, and food, and wild individuals often bear scars from previous conflicts ^8^.

To understand the genetic underpinnings of the specialized Komodo dragon physiology, we sequenced its genome and used comparative genomics to discover how the Komodo dragon genome differs from other species. We present a high quality *de novo* assembly, generated with a hybrid approach of long-range sequencing using 10x Genomics, PacBio, and Oxford Nanopore sequencing, and single-molecule physical mapping using the BioNano platform. This suite of technologies allowed us to confidently assemble a high-quality reference genome for the Komodo dragon, which can serve as a template for other varanid lizards. We used this genome to understand the relationship of varanids to other reptiles using a phylogenomics approach. We uncovered Komodo-specific positive selection for a suite of genes encoding regulators of muscle metabolism, cardiovascular homeostasis, and thrombosis. Further, we discovered multiple lineage-specific expansions of a family of chemoreceptor genes in several squamates, including some lizards and a snake, as well as in the Komodo dragon genome. Finally, we generated a high-resolution chromosomal map resulting from assignment of scaffold to chromosomes, providing a powerful tool to address questions about karyotype and sex chromosome evolution in squamates.

## Results

### De novo genome assembly

We obtained DNA from peripheral blood of two male individuals housed at Zoo Atlanta: Slasher, a male offspring of the first Komodo dragons given as gifts in 1986 to US President Reagan from President Suharto of Indonesia, and Rinca, a juvenile male (Figure 1A). The *V. komodoensis* genome is distributed across 20 pairs of chromosomes, comprising eight pairs of large chromosomes and 12 pairs of microchromosomes ^13,14^. *De novo* assembly was performed using 57X coverage of 10x Genomics linked-read sequencing data to generate an initial assembly with a scaffold N50 of 10.2 Mb and a contig N50 of 95 kb. Separately, 80X coverage of Bionano physical mapping data was *de novo* assembled to create an assembly with a scaffold N50 of 1.2 Mb. These two assemblies were merged into a hybrid assembly and then scaffolded further using 6.3X coverage from PacBio sequencing and 0.75X coverage from Oxford Nanopore MinIon sequencing, for a total coverage of 144X. The final assembly contained 1,403 scaffolds (>10 kb) with an N50 of 29 Mb (longest scaffold: 138 Mb) (Table 1). The assembly is 1.51 Gb in size, ~32% smaller than the genome of the Chinese crocodile lizard (*Shinisaurus crocodilurus*), the closest relative of the Komodo dragon for which a sequenced genome is available, and ~15% smaller than the green anole (*Anolis carolinensis*), a model squamate lizard (Table S1). An assembly-free error corrected *k-*mer counting estimate of genome size estimates the Komodo dragon genome to be 1.69 Gb, making our assembly 89% complete. The GC content of the Komodo dragon genome is 44.0%, similar to the GC content of the *S. crocodilurus* genome (44.5%) but higher than the GC content of *A. carolinensis* (40.3%) (Table S1). Repeats were annotated using RepeatMasker and the squamate repeat dabatase ^15^. Repetitive elements accounted for 32% of the genome, most of which were transposable elements (Table S2). As repetitive elements in *S. crocodilurus* account for 49.6% of the genome ^16^, most of the difference in size between the Komodo dragon genome and its closest sequenced relative genome can be attributed to differences in repetitive element content.

**Table 1.** Genome statistics of the Komodo dragon genome.

|  |  |
| --- | --- |
| Assembly size | 1.51 Gb (1,508,391,850 bp) |
| Number of scaffolds | 1,403 |
| Minimum scaffold length | 10 Kb |
| Maximum scaffold length | 138 Mb |
| N50 | 29 Mb (29,129,838) |
| Number of protein-coding genes | 18,462 |
| GC content | 44.04% |

**Figure 1.**
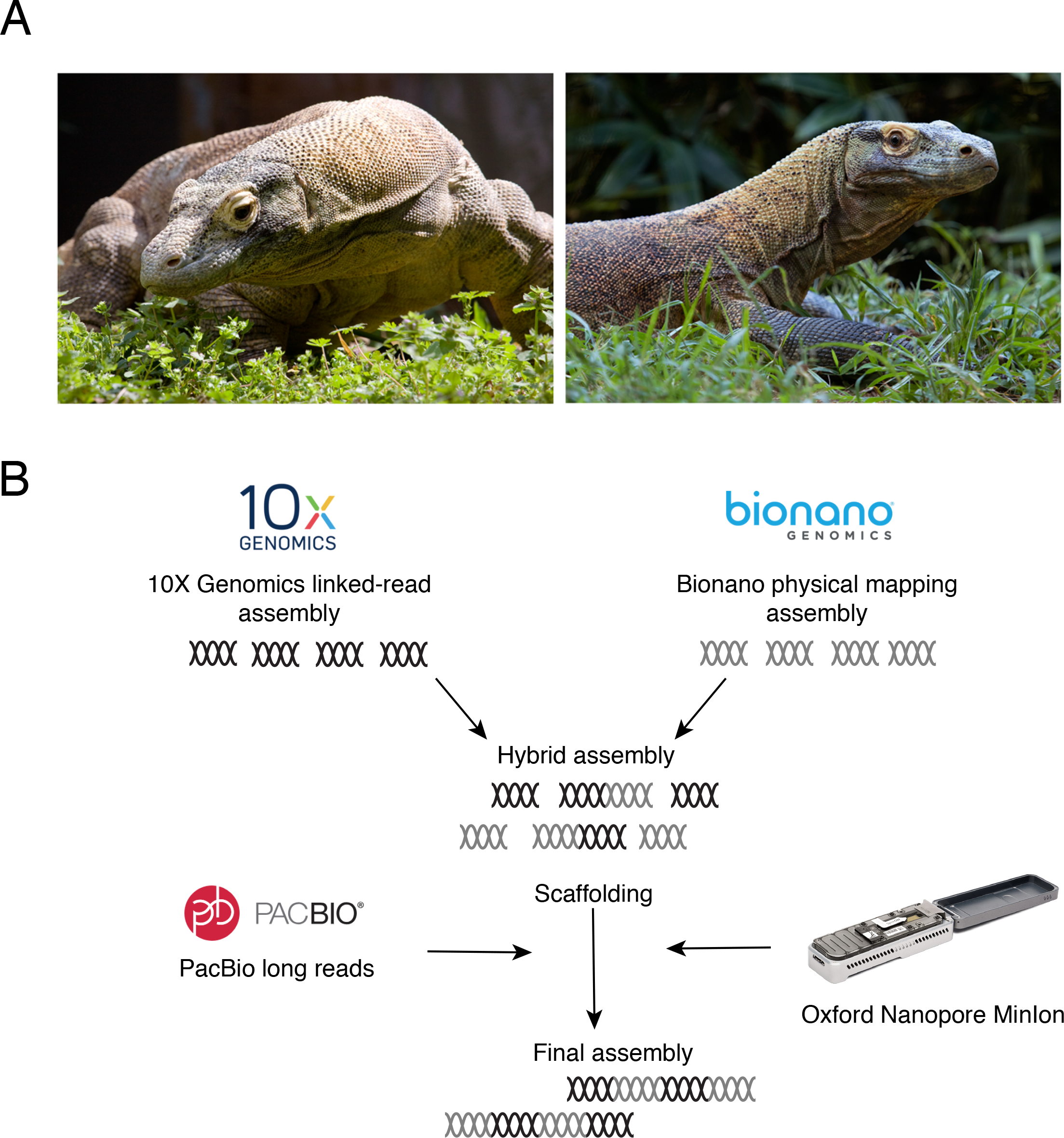
(A) Komodo dragons (left, Slasher; right, Rinca) sampled for DNA in this study. Photos courtesy of Adam K Thompson/Zoo Atlanta. (B) Genome assembly workflow. Two separate *de novo* assemblies were generated with 10x genomics and Bionano physical mapping data and merged into an intermediate hybrid assembly. Long reads from PacBio and Oxford Nanopore MinIon were used to scaffold the hybrid assembly into a final version.

### Chromosome scaffold content

We isolated chromosome-specific DNA pools from a female embryo of *V. komodoensis* (VKO) from Prague zoo stock through flow sorting ^14^. We obtained Illumina short-read sequences of these 15 DNA pools containing all chromosomes of *V. komodoensis* (Table S3). For each chromosome we determined scaffold content and homology to *A. carolinensis* and *G. gallus* chromosomes (Table 2 and Table S4). For each chromosome, we determined scaffold content and homology to *A. carolinensis* and *G. gallus* chromosomes (Table 2 and Tables S4-5). For those pools where chromosomes were mixed (VKO6/7, VKO8/7, VKO9/10, VKO11/12/W, VKO17/18/Z, VKO17/18/19) we determined partial scaffold content of single chromosomes. Homology to *A. carolinensis* microchromosomes was determined using scaffold assignments to chromosomes performed in *Anolis* chromosome-specific sequencing project ^17^. A total of 243 scaffolds containing 1.14 Gb (75% of total 1.51 Gb assembly) were assigned to 20 chromosomes of *V. komodoensis*. As sex chromosomes shares homologous regions (pseudoautosomal regions), scaffolds that were enriched in both 17/18/Z and 11/12/W samples most likely contained sex chromosome regions of *V. komodoensis*. Considering that our reference genome was from a male individual, they were assigned to the Z chromosome. All these scaffolds were homologous to *A. carolinensis* chromosome 18, and mostly to chicken chromosome 28 as recently determined by transcriptome analysis ^18^.

**Table 2.**
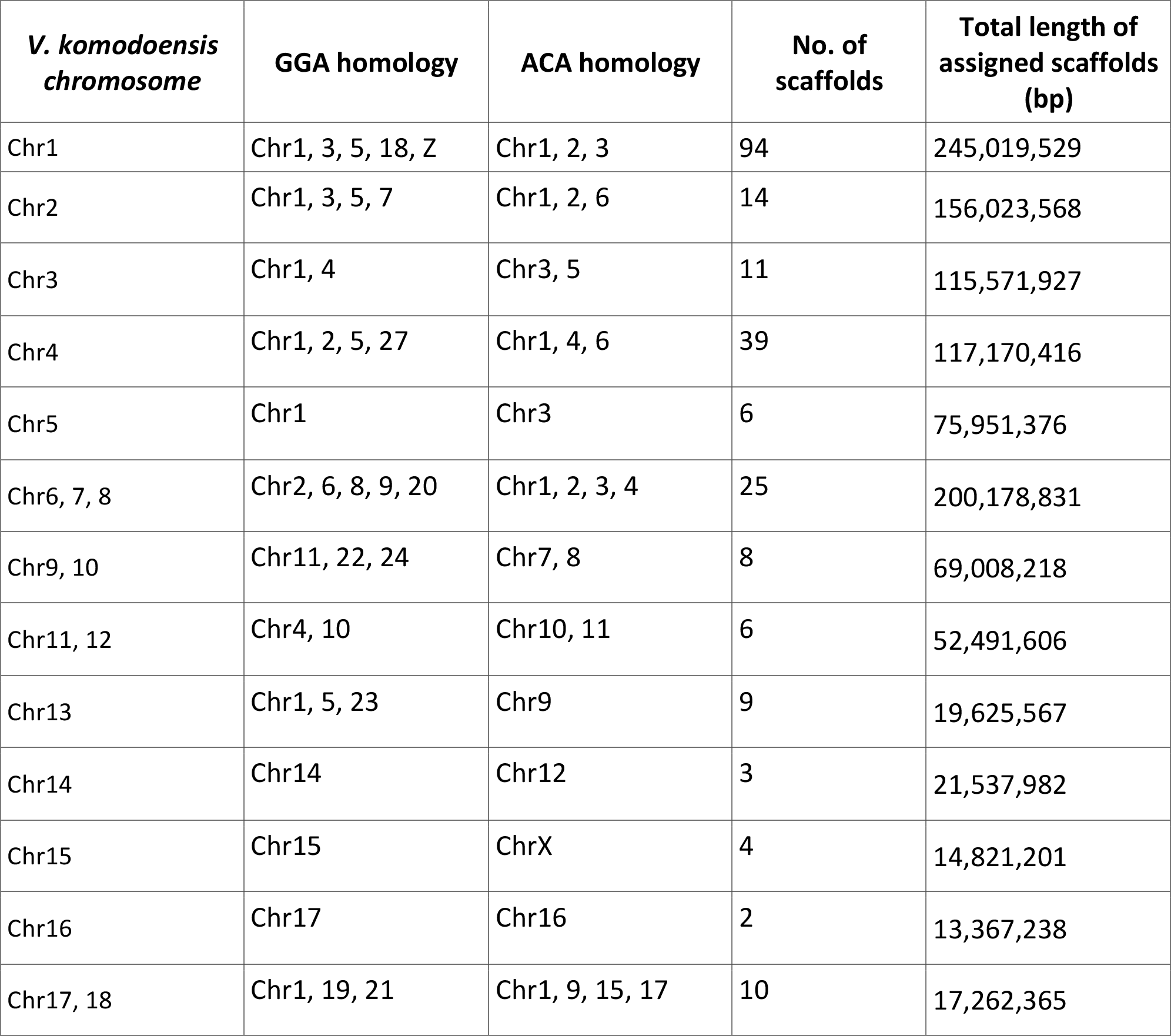
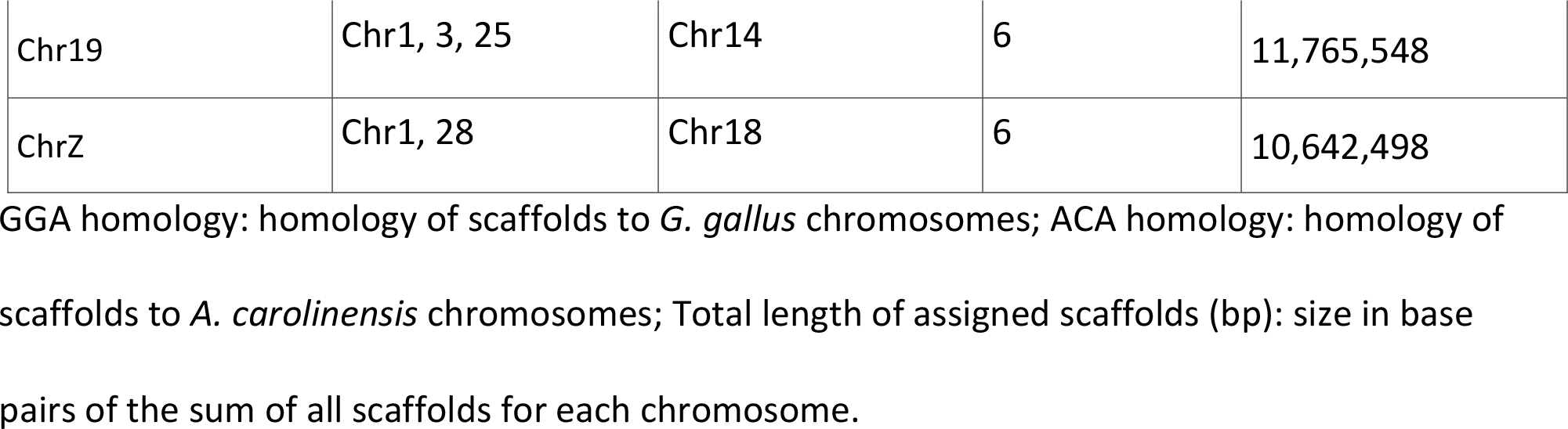
Results of scaffold assignments to chromosomes of *V. komodoensis*.

### Gene annotation

As Komodo dragons have a unique cardiovascular physiology, we used heart tissue as the source for RNA sequencing to increase the accuracy of cardiovascular gene prediction, increasing our power to detect interesting changes to the cardiovascular system encoded in the genome. RNA sequencing was assembled into transcripts with Trinity ^19^. After soft masking repetitive elements, genes were annotated using the MAKER pipeline with protein homology, assembled transcripts, and de novo predictions as evidence, and stringently quality filtered (see Methods). A total of 18,462 protein coding genes were annotated in the Komodo genome, 17,194 (93%) of which have one or more annotated Interpro functional domain (Table 1). Of these protein-coding genes, 63% were expressed (RPKM > 1) in the heart. A total of 89% of Komodo dragon protein-coding genes are orthologous to genes in the model lizard *A. carolinensis* genome. The median percent identity of single-copy orthologs between Komodo and *A. carolinensis* is 68.9%, whereas it is 70.6% between single-copy orthologs in Komodo and *S. crocodilurus* (Figure S1).

### Phylogenetic placement of Komodo

As the Komodo dragon genome is the first monitor lizard (Family Varanidae) to have a complete genome sequence, previous phylogenetic analyses of varanid lizards has been limited to marker sequences ^20,21^. We used the Komodo dragon genome to estimate a species tree using 2,752 single copy orthologs (see Methods) present in the Komodo dragon and 14 representative non-avian reptile species, including 7 squamates, 3 turtles, and 4 crocodilians, along with one avian species (chicken) and one mammalian species (mouse) (Figure 2). The placement of Komodo dragon and the monitor lizard genus using this genome-wide dataset agrees with previous marker gene studies ^20,21^.

**Figure 2.**
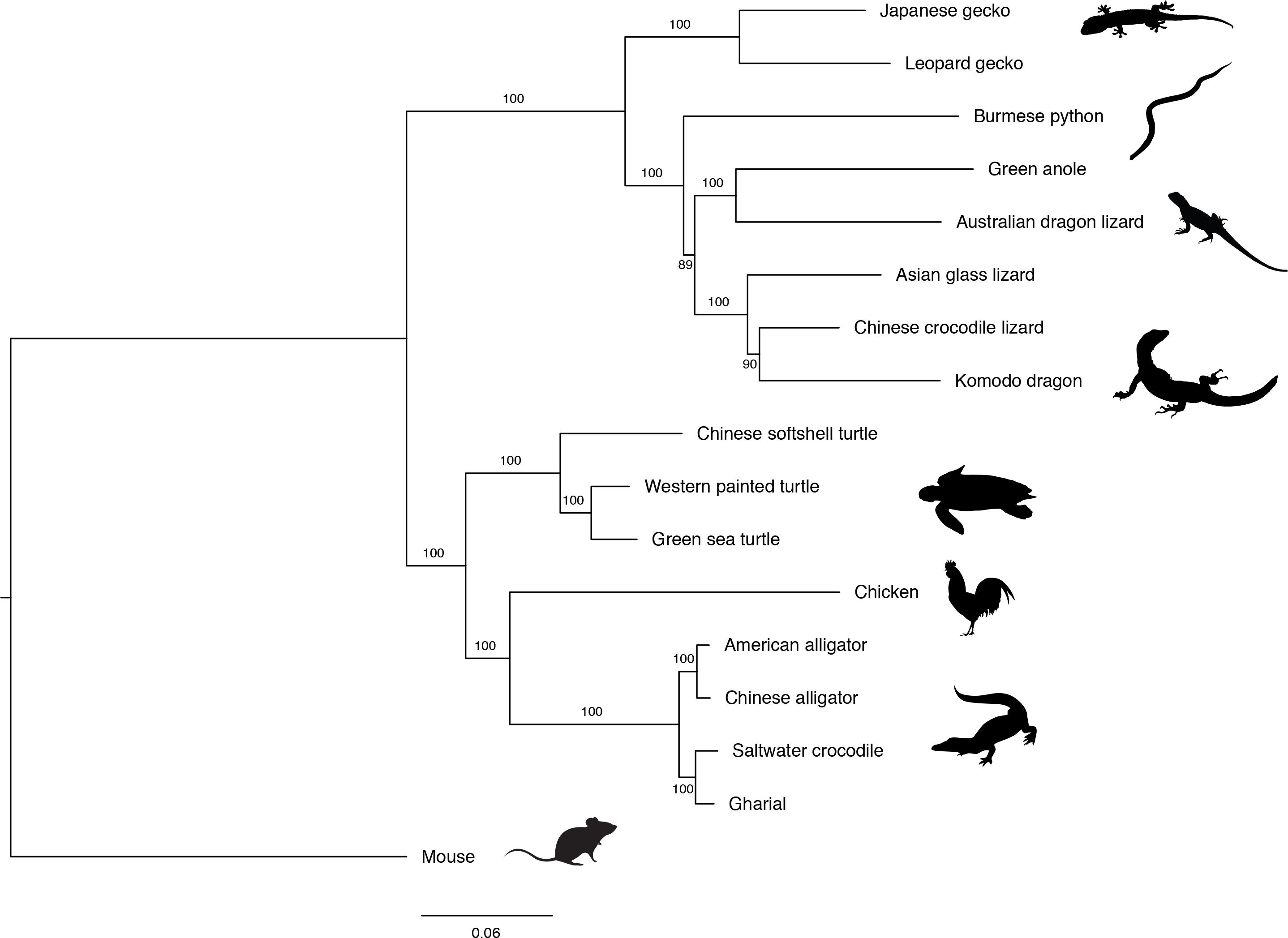
Estimated species phylogeny of 15 non-avian reptiles species and 2 additional vertebrates. Maximum likelihood phylogeny was constructed from 2,752 single-copy orthologous proteins. Support values from 10,000 bootstrap replicates are shown. All images obtained from PhyloPic.org.

### Expansion of vomeronasal genes across squamate reptiles

The vomeronasal, or the Jacobson’s, organ is a chemosensory tissue that detects chemical cues such as pheromones and kairomones. It is shared across amphibians, mammals, and reptiles though it has been secondarily lost in some groups, including birds ^22,23^. Squamate reptiles such as snakes and lizards have apparently functional vomeronasal organs with the ability to sense prey-derived chemical signals, as well as specific associated behaviors such as tongue-flicking to deliver olfactory cues to the sensory tissue, and it is clear that the vomeronasal organ plays an important role in squamate reptile ecology ^24^. Two types of chemosensory receptors, both of which are seven-transmembrane G-protein coupled receptors, function as sensors in the vomeronasal organ. The number of Type 1 vomeronasal receptors (V1Rs) has expanded through gene duplications in certain mammalian lineages, while the number of Type 2 receptors (V2Rs) has expanded in amphibians and some mammalian lineages ^22^. Crocodilian and turtle genomes contain few to no V1R and V2R genes ^25^. Snakes, in contrast, have a significantly expanded V2R repertoire that has arisen through gene duplication^26^.

To clarify the relationship between vomeronasal organ function and evolution of vomeronasal-receptor gene families, we analyzed the coding sequences of 15 reptiles, including Komodo, for presence of V1R and V2R genes (Figure 3A). We confirmed that there are few V1R genes across reptiles generally and few to no V2R genes in crocodilians and turtles (Table S6). The low number of V2R genes in green anole (*Anolis carolinensis)* and Australian dragon lizard (*Pogona vitticeps)* suggest that V2R genes are infrequently expanded in iguanians, though more iguanian genomes are needed to test this hypothesis. In contrast, we found a large repertoire of V2Rs, comparable in size to that of snakes, in the Komodo dragon and other lizards.

**Figure 3.**
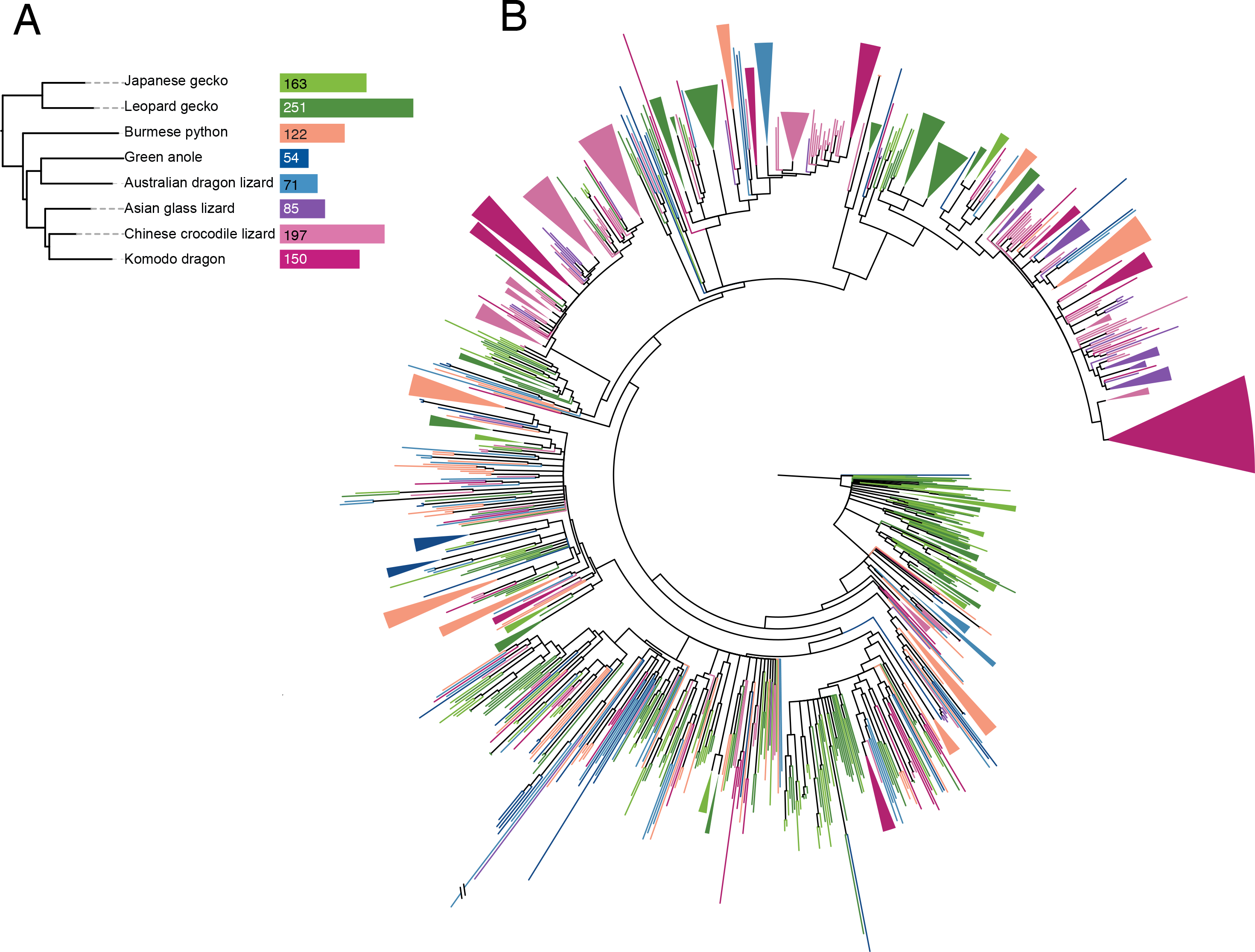
Type 2 vomeronasal receptors have expanded in Komodo dragons and several other squamate reptiles. (A) Type 2 vomeronasal gene counts in squamate reptiles. (B) Unrooted gene phylogeny of 1,093 vomeronasal Type 2 receptor transmembrane domains across squamate reptiles. The topology of the tree supports a gene expansion ancestral to squamates (i.e., clades containing representatives from all species) as well as multiple species-specific expansions through gene duplication events (i.e., clades containing multiple genes from one species). Branches with bootstrap support less than 60 are collapsed. Colors correspond to species in (A). Clades containing genes from a single species are collapsed.

To infer the details of the dynamic evolution of this gene family, we built a phylogeny of all V2R gene sequences across squamates (Figure 3B). The topology of this phylogeny supports that, as previously hypothesized, V2Rs expanded in the common ancestor of squamates, as there are clades of gene sequences containing members from all species ^26^. In addition, there are a large number of well-supported single species clades (i.e., Komodo dragon only) dispersed across the gene tree, which is consistent with multiple duplications of V2R genes later in squamate evolution, including in the Komodo and gecko lineages (Figure 3B).

Because V2Rs have expanded in rodents through tandem gene duplications that produced clusters of paralogs ^27^, we examined clustering of V2R genes in our Komodo assembly to determine if a similar mechanism is likely driving these gene expansions. Of 151 V2Rs, 99 are organized into 26 gene clusters ranging in size from 2 to 14 genes (Figure 4A, Table S7). To understand if these gene clusters arose through tandem gene duplication, we constructed a phylogeny of all Komodo dragon V2R genes (Figure 4B). The largest V2R cluster contains 14 V2R genes, which group together in a gene tree of Komodo V2R genes (Figure 4). Of the remaining 52 V2R genes, 38 are on scaffolds less than 10 Kb in size, so our estimate of V2R clustering is a lower bound due to fragmentation in the genome assembly (Table S7).

**Figure 4.**
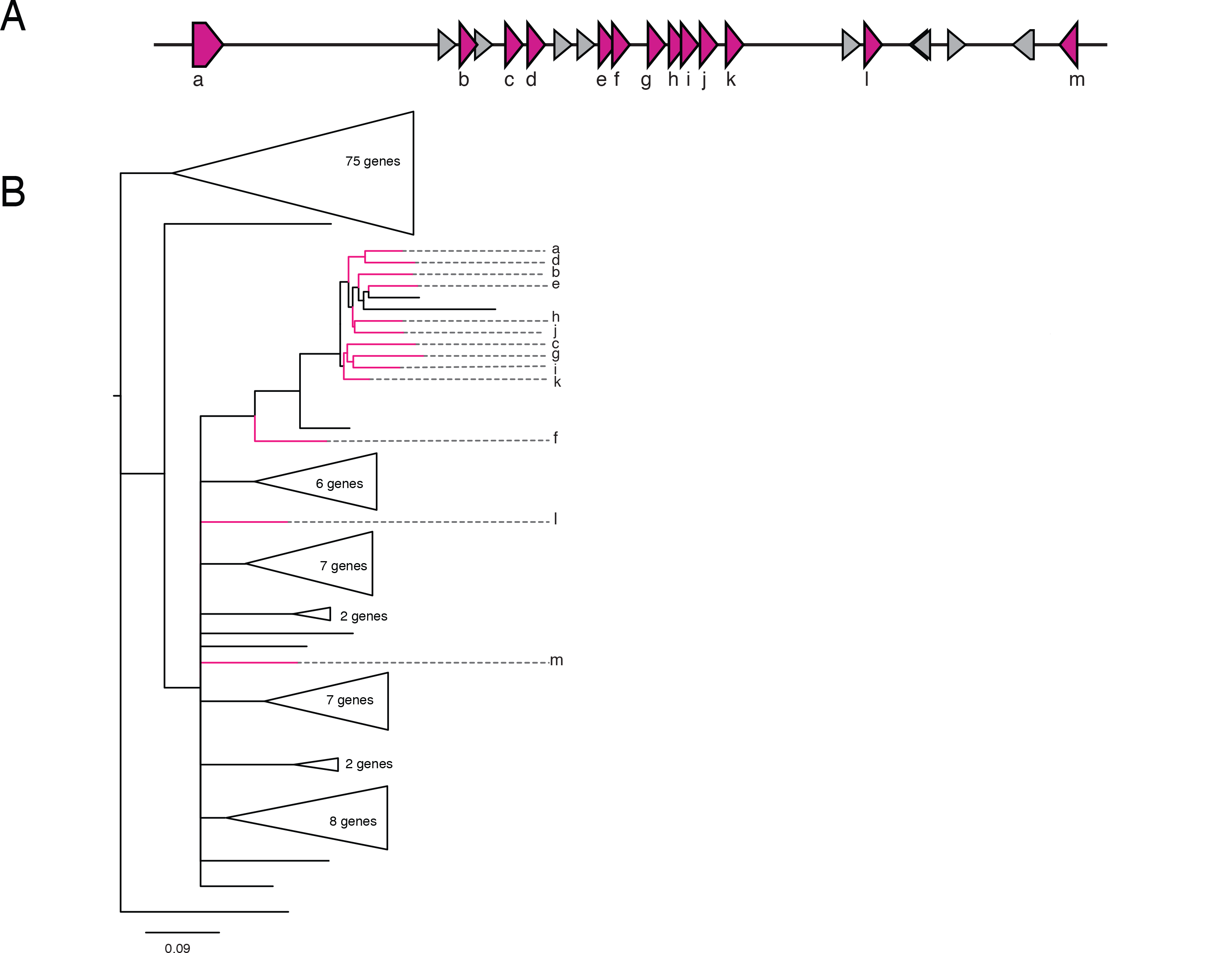
Gene clusters of Type 2 vomeronasal receptors evolved through gene duplication. (A) Genes in a cluster of vomeronasal Type 2 receptor genes in the Komodo dragon genome containing 14 V2R genes. Pink genes are V2R genes and gray genes are non-V2R genes. Gene labels correspond to labels in (B). (B) Unrooted phylogeny of 151 vomeronasal Type 2 receptor genes in Komodo. As most of the genes in this gene cluster group together in a gene phylogeny of all Komodo dragon V2R genes, it is likely that this cluster evolved through gene duplication events. Branches with bootstrap support less than 80 are collapsed. Clades without genes in this V2R gene cluster are collapsed. Genes in the V2R cluster are colored pink and labeled as in (A).

### Positive selection

To test for adaptive protein evolution in the Komodo dragon genome, we identified single-copy orthologs across squamate reptiles, built codon alignments, and ran tests of positive selection using a branch-site model to determine genes that have diversified in the varanid lineage (see Methods and Table S8). Our analysis revealed 201 genes with signatures of positive selection in Komodo dragons (Table S9). Many of the genes under positive selection point towards important adaptations of the Komodo dragon’s mammalian-like cardiovascular and metabolic functions, which are unique among non-varanid ectothermic reptiles, though 25% of positively selected genes were not detectably expressed in the heart and likely represent adaptations in other aspects of Komodo dragon biology. Pathways with positively expressed genes include mitochondrial regulation and cellular respiration, hemostasis and the coagulation cascade, innate and adaptive immunity, and angiotensinogen (a central regulator of cardiovascular physiology). Many of these have implications for Komodo physiology, and for varanid lizard physiology generally. We identified several functional categories with multiple positively selected for more detailed analysis. In each case, the genes are located in different parts of the Komodo genome and therefore likely represent recurrent selection on these functions during Komodo evolution.

### Positive selection of genes regulating mitochondrial function

Mitochondria regulate energy production in cells through the oxidative phosphorylation process, which is mediated through the electron transport chain. Multiple subunits and assembly factors of the Type 1 NADH dehydrogenase and cytochrome c oxidase protein complexes, which perform the first and last steps of the electron transport chain respectively, show evidence of positive selection in the Komodo dragon genome (Figure 5, Figure S2, Table S9). These include the genes *NDUFA7*, *NDUFAF7*, *NDUFAF2*, *NDUFB5* from the Type 1 NADH dehydrogenase complex and *COX6C* and *COA5* from the cytochrome c oxidase complex.

**Figure 5.**
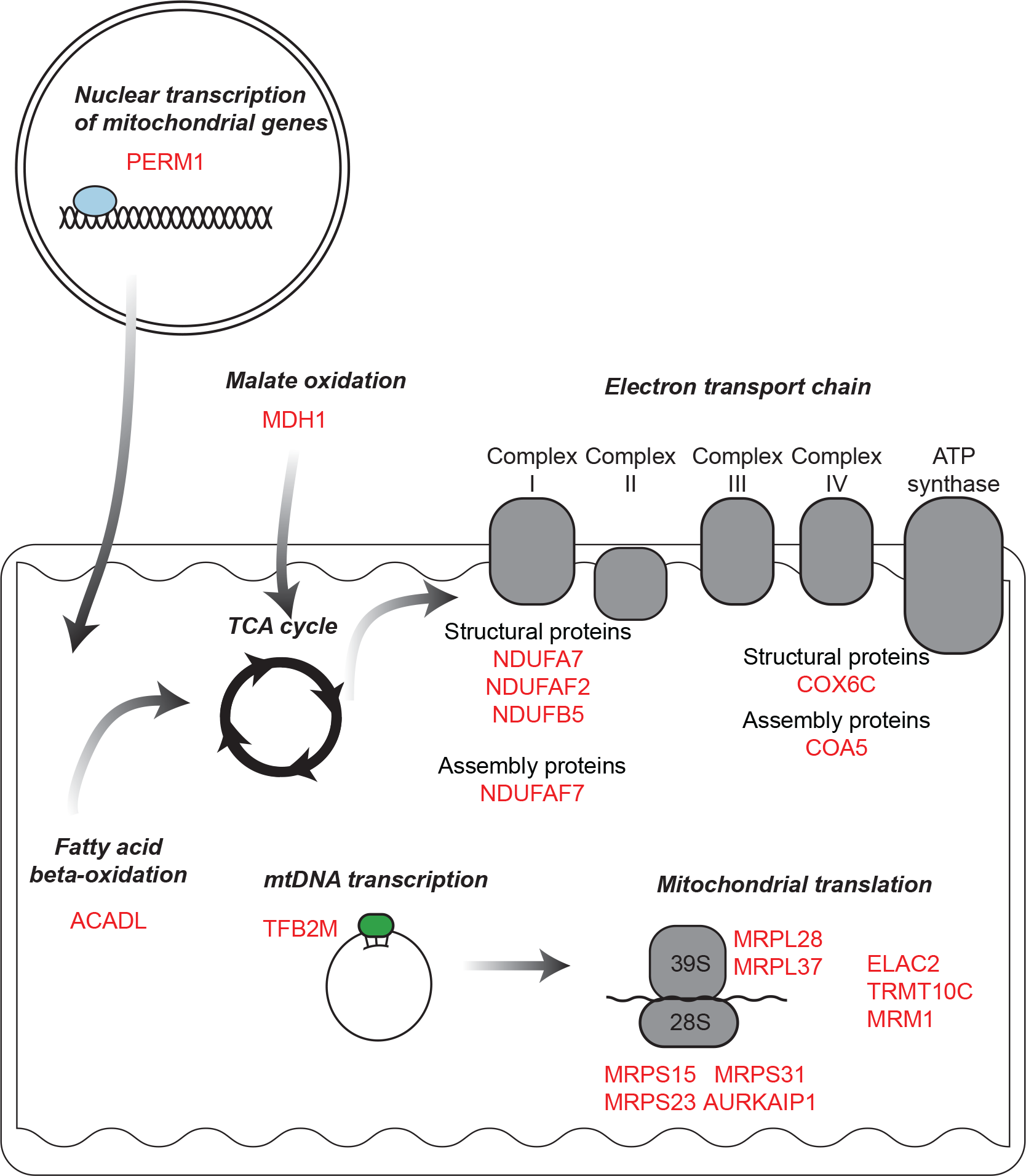
Mitochondrial genes under positive selection in the Komodo dragon. Genes in the Komodo dragon genome under positive selection include components of the electron transport chain, regulators of transcription, regulators of translation, and fatty acid beta-oxidation.

Beyond the electron transport chain, other elements of mitochondrial function have signatures of positive selection in the Komodo lineage (Figure 5). Of note, we also detected positive selection for *ACADL*, which encodes LCAD - acyl-CoA dehydrogenase, long chain—a member of the acyl-CoA dehydrogenase family. LCAD is a critical enzyme for mitochondrial fatty acid beta-oxidation, the major postnatal metabolic process in cardiac myocytes ^28^. Further, two genes that promote mitochondrial biogenesis, *TFB2M* and *PERM1*, have undergone positive selection in the Komodo dragon. *TFB2M* regulates mtDNA transcription and dimethylates mitochondrial 12s rRNA ^29,30^. *PERM1* regulates the expression of selective PPARGC1A/B and ESRRA/B/G target genes with roles in glucose and lipid metabolism, energy transfer, contractile function, muscle mitochondrial biogenesis and oxidative capacity ^31^. *PERM1* also enhances mitochondrial biogenesis, oxidative capacity, and fatigue resistance when over-expressed in mice ^32^. Finally, we also identified *MDH1*, encoding malate dehydrogenase, which together with the mitochondrial *MDH2*, regulates the supply of NADPH and acetyl-CoA to the cytoplasm, thus modulating fatty acid synthesis ^33^.

Multiple factors regulating translation within the mitochondria have also undergone positive selection in the Komodo dragon (Figure 5). This includes the mitochondrial ribosome, including four components of 28S small ribosomal subunit (*MRPS15*, *MRPS23*, *MRPS31*, and *AURKAIP1*) and two components of the 39S large ribosomal subunit (*MRPL28* and *MRPL37*). We also found evidence for positive selection on the *ELAC2* and *TRMT10C* genes, which are required for maturation of mitochondrial tRNA, and MRM1, which encodes a mitochondrial rRNA methyltransferase ^34–36^.

Overall, these instances of positive selection in a large range of genes encoding proteins important for mitochondrial function and biogenesis clearly point to a coordinated genetic pathway that could explain the remarkable aerobic capacity of the Komodo dragon. While it is not possible to determine whether these adaptations are present in other monitor lizards in the absence of a sequenced genome, changes in cellular respiration likely play a role in the high aerobic capacity of most varanid lizards.

### Positive selection of angiotensinogen

We detected positive selection for angiotensinogen (*AGT*), which encodes the precursor of several important peptide regulators of cardiovascular function, the most well-studied being angiotensin II (AII) and angiotensin1-7 (A1-7). AII has multiple important and potent activities in cardiovascular physiology. The two most notable, and perhaps most relevant to Komodo dragon physiology, are its vasoactive function in blood vessels, and its inotropic effects on the heart. In mammals during intense physical activity, All increases and contributes to arterial blood pressure and regional blood regulation ^37,38^. The positive selection for *AGT* points to important adaptations in these physiological parameters. Reptiles have a functional renin-angiotensin system that is important for their cardiovascular physiology ^39–41^. It is likely that positive selection for *AGT* is related to a mammalian-like cardiovascular function in the Komodo dragon.

### Positive selection of thrombosis-related genes

We find evidence for positive selection across different elements of the coagulation cascade, including regulators of platelets and fibrin. The coagulation cascade controls thrombosis, or blood clotting, preventing blood loss during injury. Four genes that regulate platelet activities, *MRVI1*, *RASGRP1*, *LCP2*, and *CD63* have undergone positive selection in the Komodo dragon genome. *MRVI1* is involved in inhibiting platelet aggregation ^42^, *RASGRP1* coordinates calcium dependent platelet responses ^43^, *LCP2* is involved in platelet activation ^44^, and *CD63* plays a role in controlling platelet spreading ^45^. In addition to regulators of platelets, two coagulation factors, *F10* (Factor X) and *F13B* (Coagulation factor XIII B chain) have undergone positive selection in the Komodo genome. Factor X is centrally important to the coagulation cascade and its activation is the first step in initiating coagulation ^46^. Factor 13B is the beta subunit of Factor 13, which is the final coagulation factor activated in the coagulation cascade ^47^. Further, *FGB*, which encodes one of the three subunits of fibrinogen, the molecule which is converted to the clotting agent fibrin ^48^, has undergone positive selection in the Komodo genome.

Komodo dragons, along with other species of monitor lizards, produce anticoagulants and hypotension-inducing proteins in their saliva which are hypothesized to aid in hunting ^11,12^. In addition to hunting, Komodo dragons use their serrate teeth during intraspecific conflict, which can be aggressive and inflict serious wounds ^8^. Because it is likely that their saliva enters the bloodstream of Komodo dragons during these conflicts, we hypothesize that the positive selection that we detected in many Komodo dragon coagulation genes may result from selective pressure for Komodo dragons to evade the anticoagulant effects of conspecifics.

### Discussion

We have sequenced and assembled a high-quality genome of the Komodo dragon. The combination of platforms that we used allowed the de novo assembly of a genome that will serve as a template for analysis of other varanid genomes, and for further investigation of genomic innovations in the varanid lineage. Moreover, we assigned 75% of the genome to chromosomes. Assignment of the Komodo dragon genome to chromosomes provides a significant contribution to comparative genomics of squamates and vertebrates in general.

Our comparative genomic analysis identified previously undescribed species-specific expansion of Type 2 vomeronasal receptors across multiple squamates, including lizards and at least one snake. It will be exciting to explore the role this expansion of V2Rs plays in behavior and ecology of Komodo dragons, including their ability to locate prey at long distances ^8^. Komodo dragons, like other squamates, are known to possess a sophisticated lingual-vomeronasal systems for chemical sampling of their environment ^49^. This sensory apparatus allows Komodo dragons to perceive chemicals from the environment for a variety of social and ecological activities, including kin recognition, mate choice ^50,51^, predator avoidance ^52,53^, hunting prey ^54,55^, and for locating and tracking injured or dead prey. Komodo dragons are unusual as they adopt both foraging tactics across ontogeny with smaller juveniles preferring active foraging for small prey and large adult dragons targeting larger ungulate prey via ambush predation ^10^. However, retention of a highly effective lingual-vomeronasal system across ontogeny seems likely, given the exceptional capacity for Komodo dragons of all sizes to locate injured or dead prey.

We find evidence for positive selection across many genes involved in regulating mitochondrial biogenesis, cellular respiration, and cardiovascular homeostasis. Komodo dragons, along with other monitor lizards, have a high aerobic capacity and exercise endurance, and our results reveal selective pressures on biochemical pathways that are likely to be the source of this high aerobic capacity. Future genomic work on additional varanid species, and other squamate outgroups, will test these hypotheses. These selective processes are consistent with the increased oxidative capacity in python hearts after feeding ^28^. Reptile muscle mitochondria typically oxidize substrates at a much lower rate than mammals, partly based on substrate-type use ^56^. The findings that Komodo have experienced selection for several genes encoding mitochondrial enzymes, including one involved in fatty acid metabolism, points towards a more mammalian-like mitochondrial function. In addition to a clear indication of adaptive muscle metabolism, we found positive selection for AGT, which encodes two potent vasoactive and inotropic peptides with central roles in cardiovascular physiology. A compelling hypothesis is that this positive selection is an important component in the ability of the Komodo to rapidly increase blood pressure and cardiac output for attacks on prey, extended periods of locomotion including inter-island swimming, and male-male combat during the breeding season. Direct measures of cardiac function have not been made in Komodo dragons, but in other varanid lizards, a large aerobic scope during exercise is associated with a large factorial increase in cardiac output ^57^. Overall, these cardiovascular genes suggest a profoundly different cardiovascular and metabolic profile relative to other squamates, endowing the Komodo dragon with unique physiological properties.

We also found evidence for positive selection across genes that regulate blood clotting. Like other monitor lizards, the saliva of Komodo dragons contains anticoagulants. The extensive positive selection on the genes encoding their coagulation system likely reflects that there is selective pressure for Komodo dragons to evade the anticoagulant and hypotensive effects of the saliva of conspecific rivals for food, territories, or mates. While all monitor lizards tested contain anticoagulants in their saliva, the precise mechanism by which they act varies ^12^. It is likely that monitor lizards have evolved different types of adaptations that reflect the diversity of their anticoagulants. Understanding how these systems have evolved has the potential to further our understanding of the biology of thrombosis.

Varanids, including Komodo dragons, possess genotypic sex determination and share ZZ/ZW sex chromosomes with other anguimorphan lizards ^14,18^. Here, we were able to detail the content of Z chromosome of *V. komodoensis*. The chromosome sequencing data provided significant insights into the content of *V. komodoensis* Z chromosome. All scaffolds assigned to Z chromosome were homologous to *A. carolinensis* chromosome 18 (ACA18) and to chicken chromosome 28, as confirmed by comparison of blood transcriptome between sexes ^18^. The same syntenic blocks and genes appear to be implicated in different vertebrate lineages in sex determination mechanisms ^58^. In particular, the regions of sex chromosomes that are shared by the common ancestor of varanids and several other lineages of anguimorphan lizards contain the *amh* (anti-Müllerian hormone) gene ^18^, which plays a crucial role in the testis differentiation pathway. Homologs of the *amh* gene are also strong candidates for being the sex-determining genes in several lineages of teleost fishes and in monotremes ^59–62^.

## Materials and Methods

### DNA isolation and processing for Bionano optical mapping

Komodo dragon whole blood was used to extract high molecular weight genomic DNA for genome mapping. Blood was centrifuged at 2000g for 2 minutes, plasma was removed, and the sample was stored at 4°C. 2.5μl of blood was embedded in 100μl of agarose gel plug to give ~7μg DNA/plug, using the BioRad CHEF Mammalian Genomic DNA Plug Kit (Bio-Rad Laboratories, Hercules, CA, USA). Plugs were treated with proteinase K overnight at 50°C. The plugs were then washed, melted, and then solubilized with GELase (Epicentre, Madison, WI, USA). The purified DNA was subjected to four hours of drop-dialysis. DNA concentration was determined using Qubit 2.0 Fluorometer (Life Technologies, Carlsbad, CA, USA), and the quality was assessed with pulsed-field gel electrophoresis.

The high molecular weight DNA was labeled according to commercial protocols using the IrysPrep Reagent Kit (Bionano Genomics, San Diego, CA, USA). Specifically, 300 ng of purified genomic DNA was nicked with 7 U nicking endonuclease Nb.BbvCI (NEB, Ipswich, MA, USA) at 37°C for two hours in NEB Buffer 2. The nicked DNA was labeled with a fluorescent-dUTP nucleotide analog using Taq polymerase (NEB) for one hour at 72°C. After labeling, the nicks were repaired with Taq ligase (NEB) in the presence of dNTPs. The backbone of fluorescently labeled DNA was stained with DNA stain (BioNano).

### DNA processing for 10x Genomics linked read sequencing

High molecular weight genomic DNA extraction, sample indexing, and generation of partition barcoded libraries were performed by 10x Genomics (Pleasanton, CA, USA) according to the Chromium Genome User Guide and as published previously ^64^.

### Bionano mapping and assembly

Using the Bionano Irys instrument, automated electrophoresis of the labeled DNA occurred in the nanochannel array of an IrysChip (Bionano Genomics), followed by automated imaging of the linearized DNA. The DNA backbone (outlined by YOYO-1 staining) and locations of fluorescent labels along each molecule were detected using the Irys instrument’s software. The length and set of label locations for each DNA molecule defines an individual single-molecule map. Raw Bionano single-molecule maps were de novo assembled into consensus maps using the Bionano IrysSolve assembly pipeline (version 5134) with default settings, with noise values calculated from the 10x Genomics Supernova assembly.

### 10x Genomics sequencing and assembly

The 10x Genomics barcoded library was sequenced on the Chromium machine, and the raw reads were assembled using the company’s Supernova software (version 1.0) with default parameters. Output fasta files of the phased Supernova assemblies were generated in pseudohap format.

### Merging datasets into a single assembly

Sequencing and mapping data types were merged together as follows. First, Bionano assembled contigs and the 10x Genomics assembly were combined using Bionano’s hybrid assembly tool with the -B2 -N2 options. SSPACE-LongRead (cite https://doi.org/10.1186/1471-2105-15-211) was used in series with default parameters to scaffold the hybrid assembly using PacBio reads, Nanopore reads, and unincorporated 10x Genomics Supernova scaffolds/contigs, resulting in the final assembly.

### Assignment of scaffolds to chromosomes

Isolation of *V. komodoensis* (VKO) chromosome-specific DNA pools as previously described ^14^. Briefly, fibroblast cultivation of a female *V. komodoensis* were obtained from tissue samples of an early embryo of a captive individual. Chromosomes obtained by fibroblast cultivation were sorted using a Mo-Flo (Beckman Coulter) cell sorter. Fifteen chromosome pools were sorted in total. Chromosome-specific DNA pools were then amplified and labelled by degenerate oligonucleotide primed PCR (DOP-PCR) and assigned to their respective chromosomes by hybridization of labelled probes to metaphases. *V. komodoensis* chromosome pools obtained by flow sorting were named according to chromosomes (e.g. majority of DNA of VKO6/7 belong to chromosomes 6 and 7 of *V. komodoensis*). *V. komodoensis* pools for macrochromosomes are each specific for one single pair of chromosomes, except for VKO6/7 and VKO8/7, which contain one specific chromosome pair each (pair 6 and pair 8, respectively), plus a third pair which overlaps between the two of them (pair 7). For microchromosomes, pools VKO9/10, VKO17/18/19, VKO11/12/W and VKO17/18/Z contained more than one chromosome each, while the rest are specific for one single pair of microchromosomes. The W and Z chromosomes are contained in pools VKO11/12/W and VKO17/18/Z, respectively, together with two pairs of other microchromosomes each.

Chromosome-pool specific genetic material was amplified by GenomePlex^®^ Whole Genome Amplification (WGA) Kit (Sigma) following manufacturer protocols. DNA from all 15 chromosome pools was used to prepare Illumina sequencing libraries, which were independently barcoded and sequenced 125 bp paired-end in a single Illumina Hiseq2500 lane. Reads obtained from sequencing of flow-sorting-derived chromosome-specific DNA pools were processed with the dopseq pipeline (https://github.com/lca-imcb/dopseq) ^17,65^. Illumina adapters and WGA primers were trimmed off by cutadapt v1.13 ^66^. Then, pairs of reads were aligned to the genome assembly of *V. komodoensis* using bwa mem ^67^. Reads were filtered by MAPQ ≥ 20 and length ≥ 20 bp, and aligned reads were merged into positions using pybedtools 0.7.10 ^68,69^. Reference genome regions were assigned to specific chromosomes based on distance between positions. Finally, several statistics were calculated for each scaffold. Calculated parameters included: mean pairwise distance between positions on scaffold, mean number of reads per position on scaffold, number of positions on scaffold, position representation ratio (PRR) and p-value of PRR. PRR of each scaffold was used to evaluate enrichment of given scaffold on chromosomes. PRR was calculated as ratio of positions on scaffold to positions in genome divided by ratio of scaffold length to genome length. PRRs >1 correspond to enrichment, while PRRs <1 correspond to depletion. As the PRR value is distributed lognormally, we use its logarithmic form for our calculations. To filter out only statistically significant PRR values we used thresholds of logPRR >0 and its p-value <=0.01. Scaffolds with logPRR > 0 were considered enriched in the given sample. If one scaffold was enriched in several samples we chose highest PRR to assign scaffold as top sample. We also assigned homology of *V. komodoensis* genome to genomes of *Anolis carolinensis* (AnoCar2.0) and *Gallus gallus* (galGal3) generating alignment between genomes with LAST ^70^ and subsequently using chaining and netting technique ^71^. For LAST we used default scoring matrix and parameters of 400 for gap existence cost, 30 for gap extension cost and 4500 for minimum alignment score. For axtChain we used same distance matrix and default parameters for other chain-net scripts.

### RNA sequencing

RNA was extracted from heart tissue obtained from an adult male specimen that died of natural causes. Trizol reagent was used to extract RNA following manufacturer’s instructions. RNAseq libraries were produced using a NuGen RNAseq v2 and Ultralow v2 kits, and sequenced on an Illumina Nextseq 500.

### Genome annotation

RepeatMasker was used to mask repetitive elements in the Komodo dragon genome using the squamata repeat database as reference ^15^. After masking repetitive elements, protein-coding genes were annotated using the MAKER version 3.01.02 ^72^ pipeline, combining protein homology information, assembled transcript evidence, and de novo gene predictions from SNAP and Augustus version 3.3.1 ^73^. Protein homology was determined by aligning proteins from 15 reptile species (Table S10) to the Komodo dragon genome using exonerate version 2.2.0 ^74^. RNA-seq data was aligned to the Komodo genome with STAR version 2.6.0 ^75^ and assembled into 900,722 transcripts with Trinity version 2.4.0 ^19^. Protein domains were determined using InterProScan version 5.31.70 ^76^. Gene annotations from the MAKER pipeline were filtered based on the strength of evidence for each gene, the length of the predicted protein, and the presence of protein domains. Clusters of orthologous genes across 15 reptile species were determined with OrthoFinder v2.0.0 ^77^. A total of 284,107 proteins were clustered into 16,546 orthologous clusters. In total, 96.4% of Komodo genes were grouped into orthologous clusters. For estimating a species phylogeny only, protein sequences from *Mus musculus* and *Gallus gallus* were added to the orthologous clusters with OrthoFinder. tRNAs were annotated using tRNAscan-SE version 1.3.1^78^, and other non-coding RNAs were annotated using the Rfam database ^79^ and the Infernal software suite ^80^.

### Phylogenetic analysis

A total of 2,752 single-copy orthologous proteins across 15 reptile species, including *Varanus komodoensis, Shinisaurus crocodilurus, Ophisaurus gracilis, Anolis carolinensis, Pogona vitticeps, Python molorus bivittatus, Eublepharis macularius, Gekko japonicus, Pelodiscus sinensis, Chelonia mydas, Chrysemys picta bellii, Alligator sinensis, Alligator mississippiensis, Gavialis gangeticus*, and *Crocodylus porosus*, along with the chicken *Gallus gallus* and mouse *Mus musculus*, were each aligned using PRANK v.170427 ^81^ (Table S10). Aligned proteins were concatenated into a supermatrix, and a species tree was estimated using IQ-TREE version 1.6.7.1 ^82^ with model selection across each partition ^83^ and 10,000 ultra-fast bootstrap replicates ^84^.

### Gene family evolution analysis

Gene family expansion and contraction analyses were performed with CAFE v4.2 ^85^ for the squamate reptile lineage, with a constant gene birth and gene death rate assumed across all branches.

Vomeronasal type 2 receptors were first identified in all species by containing the V2R domain InterPro domain (IPR004073) ^86^. To ensure that no V2R genes were missed, all proteins were aligned against a set of representative V2R genes using BLASTp ^87^ with an e-value cutoff of 1e-6 and a bitscore cutoff of 200 or greater. Any genes passing this threshold were added to the set of putative V2R genes. Transmembrane domains were identified in each putative V2R gene with TMHMM v2.0 ^88^ and discarded if they did not contain 7 transmembrane domains in the C-terminal region. Beginning at the start of the first transmembrane domain, proteins were aligned with MAFFT v7.310 (auto alignment strategy) ^89^ and trimmed with trimAL (gappyout) ^90^. A gene tree was constructed using IQ-TREE ^82–84^ with the JTT+ model of evolution with empirical base frequencies and 10 FreeRate model parameters, and 10,000 bootstrap replicates. Genes were discarded if they failed the IQ-TREE composition test.

### Positive selection analysis

We analyzed 4,081 genes that were universal and single-copy across all squamate lineages tested (*Varanus komodoensis, Shinisaurus crocodilurus, Ophisaurus gracilis, Anolis carolinensis, Pogona vitticeps, Python molorus bivittatus, Eublepharis macularius*, and *Gekko japonicus*) to test for positive selection (Table S8). An additional 2,040 genes that were universal and single-copy across a subset of squamate species (*Varanus komodoensis, Anolis carolinensis, Python molurus bivittatus*, and *Gekko japonicus*) were also analyzed (Table S8). We excluded multi-copy genes from all positive selection analyses to avoid confounding from incorrect paralogy inference. Proteins were aligned using PRANK ^81^ and codon alignments were generated using PAL2NAL ^91^.

Positive selection analyses were performed with the branch-site model aBSREL using the HYPHY framework ^92,93^. For the 4,081 genes that were single-copy across all squamate lineages, the full species phylogeny of squamates was used. For the 2,040 genes that were universal and single-copy across a subset of species, a pruned tree containing only those taxa was used. We discarded genes with unreasonably high dN/dS values across a small proportion of sites, as those were false positives driven by low quality gene annotation in one or more taxa in the alignment. We used a cutoff of dN/dS of less than 50 across 5% or more of sites, and a p-value of less than 0.05 at the Komodo node. Each gene was first tested for positive selection only on the Komodo branch. Genes undergoing positive selection in the Komodo lineage were then tested for positive selection at all nodes in the phylogeny. This resulted in 201 genes being under positive selection in the Komodo lineage (Table S9).

## Acknowledgements

Special thanks from B.G.B. to John Romano for inspiration and historical information. We are grateful to staff at Zoo Atlanta for care of Slasher and Rinca and help obtaining samples, Jim Pether from Reptilandia zoo in Gran Canaria in the Canary Islands, for additional samples, and R. Chadwick and N. Carli (Gladstone Genomics Core) for DNA and RNA-seq library preparation. We also thank Kristina Giorda, Rabeea Abbas, Deanna Church (10x Genomics) for 10x Genomics Chromium sequencing and Supernova assembly. This work was supported by institutional funding from the Gladstone Institutes to B.G.B and K.S.P.; an NHLBI grant to K.S.P and B.G.B (HL098179); the Younger Family gift to B.G.B.; an NHGRI grant to P.-Y.K. (R01 HG005946); NIH training grants (T32 AR007175) to A.C.Y.M and (T32 HL007731) to Y.M. M.A. and M.R. were supported by GACR 17-22141Y, M.R. was additionally supported by Charles University projects PRIMUS/SCI/46 and Research Centre (204069).

## Author contributions

A.L.L. did genome annotation and all comparative genomics analyses. Y.Y.Y.L. led the sequencing and assembly efforts with Y.M. and A.C.Y.M. A.I. sequenced isolated chromosomes with M.R under supervision of L.K., and assigned sequences with A.M., I.K, and V.T. M.F. and V.O. contributed to genome assembly the genome in the lab of R.F. with C.C. A.K.H. led the initial development of the project. W.L.E. initially assembled the transcriptomes and annotated the genome. O.A.R. provided frozen tissue samples. J.M. collected specimen blood. M.M. and M.F. isolated samples and obtained PacBio sequence in the lab of T.P. and C.C.E.S. performed PacBio sequencing. J.W.H., J.M., and C.C. provided direction on Varanid physiology. T.J. and C.C. provided direction on Komodo dragon ecology. P.-Y.K. coordinated the genomics efforts. K.S.P. directed comparative genomic analysis. B.G.B. initiated and coordinated the project. A.L., K.S.P. and B.G.B. wrote the paper with input from all authors.

---

**Figure S1.**
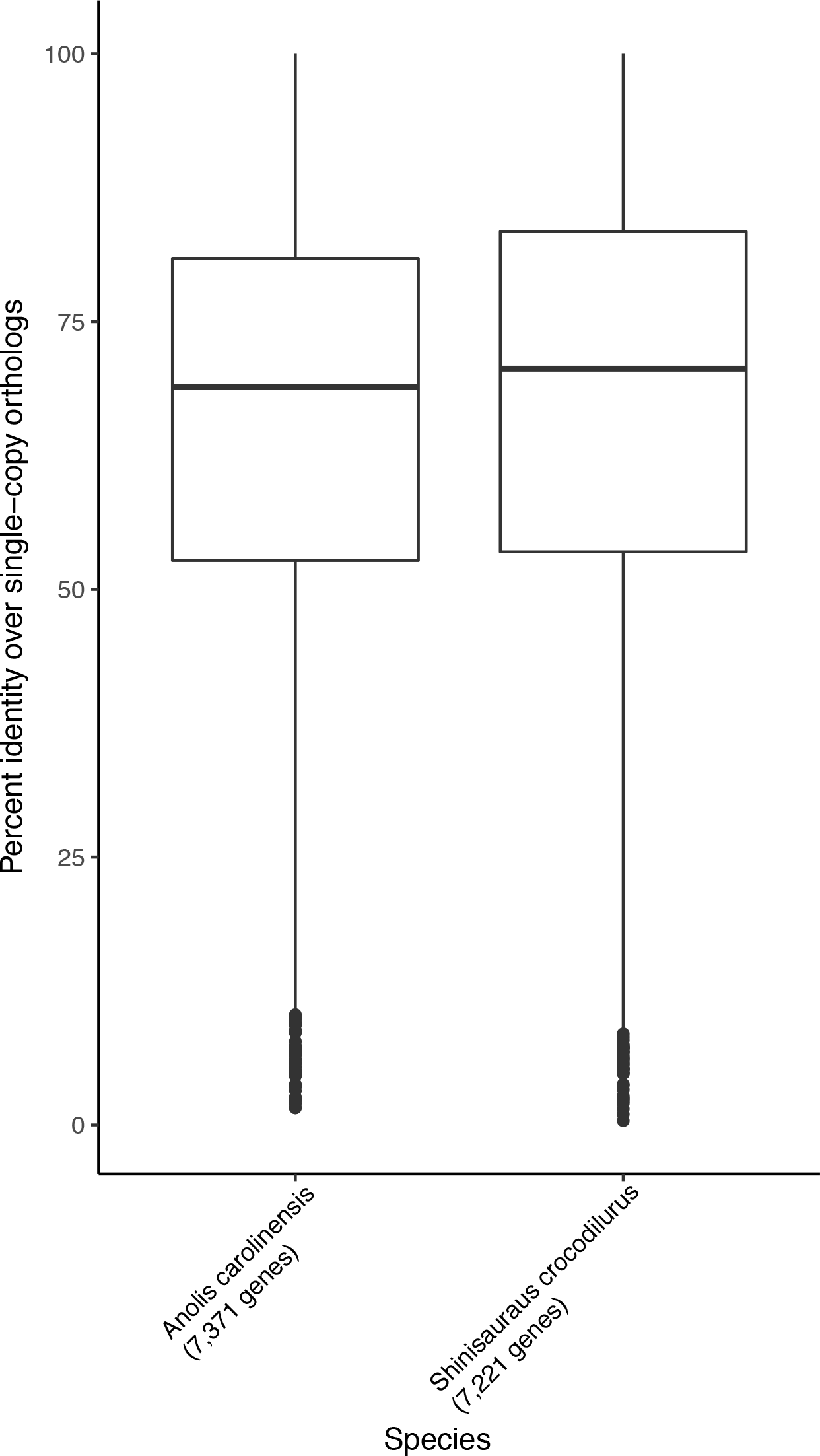
Percent identities of single-copy orthologs between the Komodo dragon and the green anole and the Komodo dragon and the Chinese crocodile lizard.

**Figure S2.**
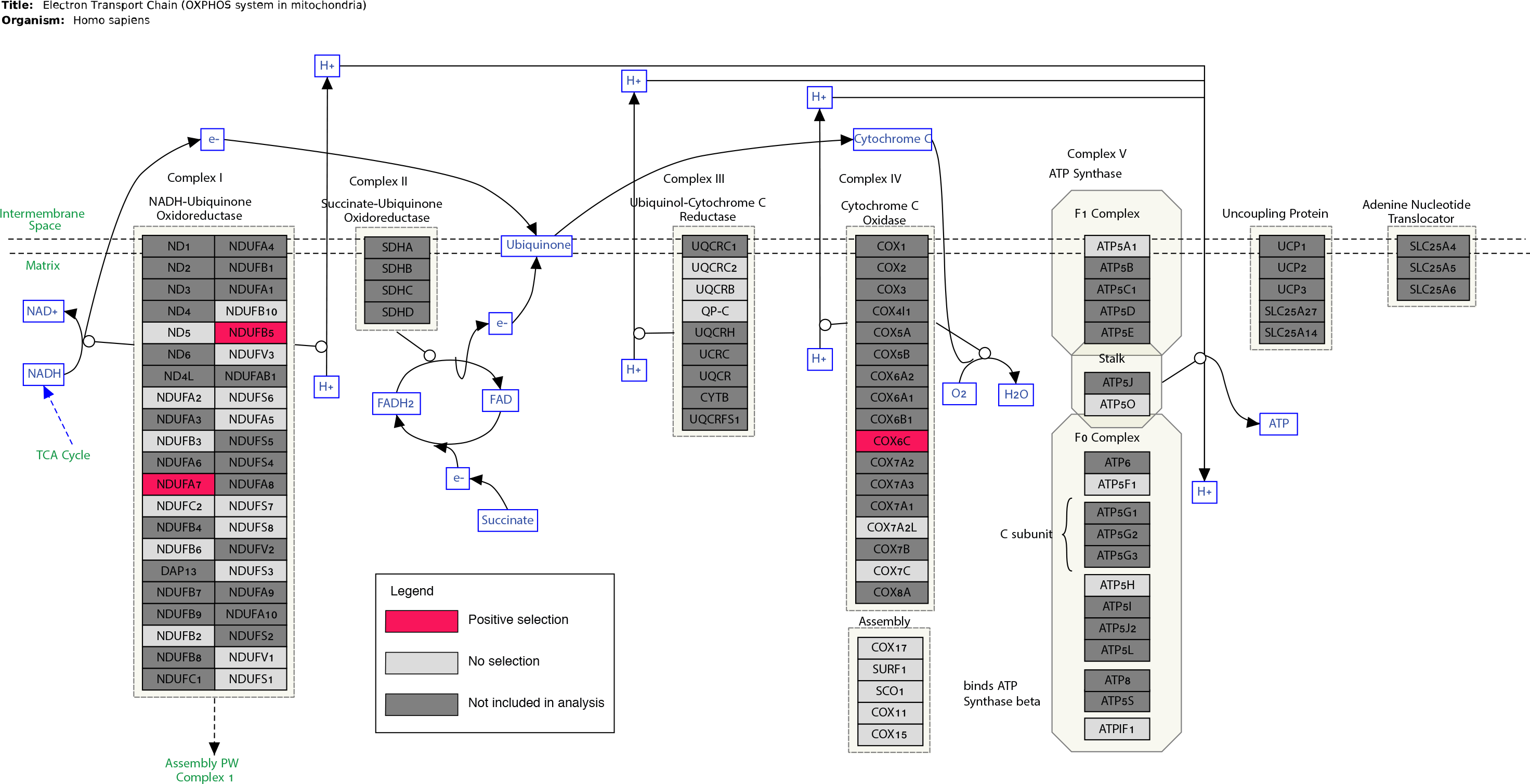
Positive selection on genes encoding structural proteins in the electron transport chain. Dark gray genes were not tested for positive selection due to either missing data in one or more species or difficulty resolving ortholog/paralog relationships. Pink genes have signatures of positive selection, and light gray genes did not have signatures of positive selection. Figure modified from WikiPathways ^63^.

## Supplemental file descriptions

Table S1. Genome statistics for non-avian reptiles used in this study.

Table S2. Repetitive elements in the Komodo dragon genome.

Table S3. Read statistics for chromosomal anchoring.

Table S4. Scaffold assignment and homologies of Komodo dragon scaffolds to green anole and chicken chromosomes.

Table S5. Number of reads, scaffolds, and positions assigned to chromosomes.

Table S6. Number of V1R/V2R genes across non-avian reptiles.

Table S7. V2R gene clusters in the Komodo dragon genome.

Table S8. Genes assayed for positive selection in the Komodo dragon genome.

Table S9. Positively selected genes in the Komodo dragon genome.

Table S10. Sources and versions of genomes used for phylogenetic and comparative methods.

## References

1. Chapman, A. D. Numbers of Living Species in Australia and the World. (Australian Biological Resources Study, 2009).

2. Collar, D. C., Schulte, J. A. & Losos, J. B. Evolution of extreme body size disparity in monitor lizards (Varanus). Evolution (N. Y). 65, 2664–2680 (2011).

3. Jensen, B., Wang, T., Christoffels, V. M. & Moorman, A. F. M. Evolution and development of the building plan of the vertebrate heart. Biochim. Biophys. Acta - Mol. Cell Res. 1833, 783–794 (2013).

4. Clemente, C. J., Withers, P. C. & Thompson, G. G. Metabolic rate and endurance capacity in Australian varanid lizards (Squamata: Varanidae: Varanus). Biol. J. Linn. Soc. 97, 664–676 (2009).

5. Burggren, W. & Johansen, K. Ventricular Haemodynamics in the Monitor Lizard Varanus Exanthematicus: Pulmonary and Systemic Pressure Separation. J. Exp. Biol. 96, (1982).

6. Ishimatsu, A., Hicks, J. W. & Heisler, N. Analysis of intracardiac shunting in the lizard, Varanus niloticus: a new model based on blood oxygen levels and microsphere distribution. Respir. Physiol. 71, 83–100 (1988).

7. King, D. & Green, B. Goannas: The Biology of Varanid Lizards. (University of New South Wales, 1998).

8. Auffenberg, W. The Behavioral Ecology of the Komodo Monitor. (University Presses of Florida, 1981).

9. Green, B., King, D., Braysher, M. & Saim, A. Thermoregulation, water turnover and energetics of free-living komodo dragons, Varanus komodoensis. Comp. Biochem. Physiol. Part A Physiol. 99, 97–101 (1991).

10. Purwandana, D. et al. Ecological allometries and niche use dynamics across Komodo dragon ontogeny. Sci. Nat. 103, 27 (2016).

11. Fry, B. G. et al. A central role for venom in predation by Varanus komodoensis (Komodo Dragon) and the extinct giant Varanus (Megalania) priscus. Proc. Natl. Acad. Sci. U. S. A. 106, 8969–8974 (2009).

12. Koludarov, I. et al. Enter the Dragon: The Dynamic and Multifunctional Evolution of Anguimorpha Lizard Venoms. Toxins 9, (2017).

13. Johnson Pokorná, M. et al. First Description of the Karyotype and Sex Chromosomes in the Komodo Dragon (Varanus komodoensis). Cytogenet. Genome Res. 148, 284–291 (2016).

14. Iannucci, A. et al. Isolating Chromosomes of the Komodo Dragon: New Tools for Comparative Mapping and Sequence Assembly. Cytogenet. Genome Res. 157, 42–50 (2019).

15. Smit, A., Hubley, R. & Green, P. Repeatmasker Open-4.0. (2013). Available at: http://www.repeatmasker.org. (Accessed: 10th January 2015)

16. Gao, J. et al. Sequencing, de novo assembling, and annotating the genome of the endangered Chinese crocodile lizard Shinisaurus crocodilurus. Gigascience 6, 1–6 (2017).

17. Kichigin, I. G. et al. Evolutionary dynamics of Anolis sex chromosomes revealed by sequencing of flow sorting-derived microchromosome-specific DNA. Mol. Genet. Genomics 291, 1955–1966 (2016).

18. Rovatsos, M., Rehák, I., Velenský, P. & Kratochvíl, L. Shared ancient sex chromosomes in varanids, beaded lizards and alligator lizards. Mol. Biol. Evol. (2019). doi:10.1093/molbev/msz024

19. Haas, B. J. et al. De novo transcript sequence reconstruction from RNA-seq using the Trinity platform for reference generation and analysis. Nat. Protoc. 8, 1494–512 (2013).

20. Pyron, R., Burbrink, F. T. & Wiens, J. J. A phylogeny and revised classification of Squamata, including 4161 species of lizards and snakes. BMC Evol. Biol. 13, 93 (2013).

21. Welton, L. J., Travers, S. L., Siler, C. D. & Brown, R. M. Integrative taxonomy and phylogeny-based species delimitation of Philippine water monitor lizards (Varanus salvator Complex) with descriptions of two new cryptic species. Zootaxa 3881, 201 (2014).

22. Silva, L. & Antunes, A. Vomeronasal Receptors in Vertebrates and the Evolution of Pheromone Detection. Annu. Rev. Anim. Biosci. 5, 353–370 (2017).

23. Stoddart, D. M. The Ecology of Vertebrate Olfaction. (Springer Netherlands, 1980).

24. Mason, R. T. & Parker, M. R. Social behavior and pheromonal communication in reptiles. J. Comp. Physiol. A 196, 729–749 (2010).

25. Green, R. E. et al. Three crocodilian genomes reveal ancestral patterns of evolution among archosaurs. Science (80-.). 346, 1254449–1254449 (2014).

26. Brykczynska, U., Tzika, A. C., Rodriguez, I. & Milinkovitch, M. C. Contrasted evolution of the vomeronasal receptor repertoires in mammals and squamate reptiles. Genome Biol. Evol. 5, 389–401 (2013).

27. Yang, H., Shi, P., Zhang, Y. & Zhang, J. Composition and evolution of the V2r vomeronasal receptor gene repertoire in mice and rats. Genomics 86, 306–315 (2005).

28. Riquelme, C. A. et al. Fatty Acids Identified in the Burmese Python Promote Beneficial Cardiac Growth. Science (80-.). 334, 528 LP–531 (2011).

29. Falkenberg, M. et al. Mitochondrial transcription factors B1 and B2 activate transcription of human mtDNA. Nat. Genet. 31, 289–294 (2002).

30. Cotney, J., McKay, S. E. & Shadel, G. S. Elucidation of separate, but collaborative functions of the rRNA methyltransferase-related human mitochondrial transcription factors B1 and B2 in mitochondrial biogenesis reveals new insight into maternally inherited deafness. Hum. Mol. Genet. 18, 2670–2682 (2009).

31. Cho, Y., Hazen, B. C., Russell, A. P. & Kralli, A. Peroxisome Proliferator-activated Receptor γ Coactivator 1 (PGC-1)- and Estrogen-related Receptor (ERR)-induced Regulator in Muscle 1 (PERM1) Is a Tissue-specific Regulator of Oxidative Capacity in Skeletal Muscle Cells. J. Biol. Chem. 288, 25207–25218 (2013).

32. Cho, Y. et al. Perm1 enhances mitochondrial biogenesis, oxidative capacity, and fatigue resistance in adult skeletal muscle. FASEB J. 30, 674–687 (2016).

33. Zhao, S. et al. Regulation of Cellular Metabolism by Protein Lysine Acetylation. Science (80-.). 327, 1000–1004 (2010).

34. Brzezniak, L. K., Bijata, M., Szczesny, R. J. & Stepien, P. P. Involvement of human ELAC2 gene product in 3’ end processing of mitochondrial tRNAs. RNA Biol. 8, 616–626 (2011).

35. Holzmann, J. et al. RNase P without RNA: Identification and Functional Reconstitution of the Human Mitochondrial tRNA Processing Enzyme. Cell 135, 462–474 (2008).

36. Lee, K.-W. & Bogenhagen, D. F. Assignment of 2ʹ-O-Methyltransferases to Modification Sites on the Mammalian Mitochondrial Large Subunit 16 S Ribosomal RNA (rRNA). J. Biol. Chem. 289, 24936–24942 (2014).

37. Forrester, S. J. et al. Angiotensin II Signal Transduction: An Update on Mechanisms of Physiology and Pathophysiology. Physiol. Rev. 98, 1627–1738 (2018).

38. Kim, S. & Iwao, H. Molecular and Cellular Mechanisms of Angiotensin II-Mediated Cardiovascular and Renal Diseases. Pharmacol. Rev. 52, 11 LP–34 (2000).

39. Wilson, J. X. The Renin-Angiotensin System in Nonmammalian Vertebrates. Endocr. Rev. 5, 45–61 (1984).

40. Fournier, D., Luft, F. C., Bader, M., Ganten, D. & Andrade-Navarro, M. A. Emergence and evolution of the renin–angiotensin–aldosterone system. J. Mol. Med. 90, 495–508 (2012).

41. Mueller, C. A., Eme, J., Tate, K. B. & Crossley, D. A. Chronic captopril treatment reveals the role of ANG II in cardiovascular function of embryonic American alligators (Alligator mississippiensis). J. Comp. Physiol. B 188, 657–669 (2018).

42. Antl, M. et al. IRAG mediates NO/cGMP-dependent inhibition of platelet aggregation and thrombus formation. Blood 109, 552–559 (2007).

43. Puetz, J. & Boudreaux, M. K. Evaluation of the gene encoding calcium and diacylglycerol regulated guanine nucleotide exchange factor I (CalDAG-GEFI) in human patients with congenital qualitative platelet disorders. Platelets 23, 401–403 (2012).

44. Bezman, N. A. et al. Requirements of SLP76 tyrosines in ITAM and integrin receptor signaling and in platelet function in vivo. J. Exp. Med. 205, 1775–88 (2008).

45. Israels, S. & McMillan-Ward, E. CD63 modulates spreading and tyrosine phosphorylation of platelets on immobilized fibrinogen. Thromb. Haemost. 93, 311–318 (2005).

46. Cooper, D. N., Millar, D. S., Wacey, A., Pemberton, S. & Tuddenham, E. G. Inherited factor X deficiency: molecular genetics and pathophysiology. Thromb. Haemost. 78, 161–172 (1997).

47. Takahashi, N., Takahashi, Y. & Putnam, F. W. Primary structure of blood coagulation factor XIIIa (fibrinoligase, transglutaminase) from human placenta. Proc. Natl. Acad. Sci. 83, 8019 LP–8023 (1986).

48. Mosesson, M. W. The roles of fibrinogen and fibrin in hemostasis and thrombosis. Semin. Hematol. 29, 177—188 (1992).

49. Halpern, M. Nasal chemical senses in reptiles: structure and function. Pp 423–523 in Gans C, Crews D (eds) Biology of the Reptilia, Vol. 18, Brain. Horm. Behav. Chicago/IL Univ. Chicago Press Google Sch. (1992).

50. Martin, J. & Lopez, P. Chemoreception, symmetry and mate choice in lizards. Proc. R. Soc. B Biol. Sci. 267, 1265–1269 (2000).

51. Baeckens, S., Martín, J., García-Roa, R. & van Damme, R. Sexual selection and the chemical signal design of lacertid lizards. Zool. J. Linn. Soc. 183, 445–457 (2018).

52. van Damme, R., Bauwens, D., Thoen, C., Vanderstighelen, D. & Verheyen, R. F. Responses of Naive Lizards to Predator Chemical Cues. J. Herpetol. 29, 38 (1995).

53. van Damme, R. & Castilla, A. M. Chemosensory predator recognition in the lizard Podarcis hispanica: effects of predation pressure relaxation. J. Chem. Ecol. 22, 13–22 (1996).

54. Cooper, W. E. Correlated evolution of prey chemical discrimination with foraging, lingual morphology and vomeronasal chemoreceptor abundance in lizards. Behav. Ecol. Sociobiol. 41, 257–265 (1997).

55. Cooper, W. Tandem evolution of diet and chemosensory responses in snakes. Amphibia-Reptilia 29, 393–398 (2008).

56. Hulbert, A. J. & Else, P. L. Evolution of mammalian endothermic metabolism: mitochondrial activity and cell composition. Am. J. Physiol. Integr. Comp. Physiol. 256, R63–R69 (1989).

57. Gleeson, T. T., Mitchell, G. S. & Bennett, A. F. Cardiovascular responses to graded activity in the lizards Varanus and Iguana. Am. J. Physiol. Integr. Comp. Physiol. 239, R174–R179 (1980).

58. Marshall Graves, J. A. & Peichel, C. L. Are homologies in vertebrate sex determination due to shared ancestry or to limited options? Genome Biol. 11, 205 (2010).

59. Hattori, R. S. et al. A Y-linked anti-Müllerian hormone duplication takes over a critical role in sex determination. Proc. Natl. Acad. Sci. 109, 2955 LP–2959 (2012).

60. Cortez, D. et al. Origins and functional evolution of Y chromosomes across mammals. Nature 508, 488 (2014).

61. Bej, D. K., Miyoshi, K., Hattori, R. S., Strüssmann, C. A. & Yamamoto, Y. A Duplicated, Truncated amh Gene Is Involved in Male Sex Determination in an Old World Silverside. G3 Genes|Genomes|Genetics 7, 2489–2495 (2017).

62. Ieda, R. et al. Identification of the sex-determining locus in grass puffer (Takifugu niphobles) provides evidence for sex-chromosome turnover in a subset of Takifugu species. PLoS One 13, e0190635 (2018).

63. Slenter, D. N. et al. WikiPathways: a multifaceted pathway database bridging metabolomics to other omics research. Nucleic Acids Res. 46, D661–D667 (2018).

64. Weisenfeld, N. I., Kumar, V., Shah, P., Church, D. M. & Jaffe, D. B. Direct determination of diploid genome sequences. Genome Res. 27, 757–767 (2017).

65. Makunin, A. I. et al. Contrasting origin of B chromosomes in two cervids (Siberian roe deer and grey brocket deer) unravelled by chromosome-specific DNA sequencing. BMC Genomics 17, 618 (2016).

66. Martin, M. Cutadapt removes adapter sequences from high-throughput sequencing reads. EMBnet.journal 17, 10 (2011).

67. Li, H. Aligning sequence reads, clone sequences and assembly contigs with BWA-MEM. (2013).

68. Quinlan, A. R. & Hall, I. M. BEDTools: a flexible suite of utilities for comparing genomic features. Bioinformatics 26, 841–2 (2010).

69. Quinlan, A. R., Pedersen, B. S. & Dale, R. K. Pybedtools: a flexible Python library for manipulating genomic datasets and annotations. Bioinformatics 27, 3423–3424 (2011).

70. Kielbasa, S. M., Wan, R., Sato, K., Horton, P. & Frith, M. Adaptive seeds tame genomic sequence comparison. Genome Res. (2011). doi:10.1101/gr.113985.110

71. Kent, W. J., Baertsch, R., Hinrichs, A., Miller, W. & Haussler, D. Evolution’s cauldron: Duplication, deletion, and rearrangement in the mouse and human genomes. Proc. Natl. Acad. Sci. 100, 11484–11489 (2003).

72. Cantarel, B. L. et al. MAKER: an easy-to-use annotation pipeline designed for emerging model organism genomes. Genome Res. 18, 188–96 (2008).

73. Stanke, M. & Morgenstern, B. AUGUSTUS: a web server for gene prediction in eukaryotes that allows user-defined constraints. Nucleic Acids Res. 33, W465–7 (2005).

74. Slater, G. & Birney, E. Automated generation of heuristics for biological sequence comparison. BMC Bioinformatics 6, 31 (2005).

75. Dobin, A. et al. STAR: ultrafast universal RNA-seq aligner. Bioinformatics 29, 15–21 (2013).

76. Jones, P. et al. InterProScan 5: genome-scale protein function classification. Bioinformatics 30, 1236–1240 (2014).

77. Emms, D. M. & Kelly, S. OrthoFinder: solving fundamental biases in whole genome comparisons dramatically improves orthogroup inference accuracy. Genome Biol. 16, 157 (2015).

78. Lowe, T. M. & Eddy, S. R. tRNAscan-SE: a program for improved detection of transfer RNA genes in genomic sequence. Nucleic Acids Res. 25, 955–64 (1997).

79. Griffiths-Jones, S., Bateman, A., Marshall, M., Khanna, A. & Eddy, S. R. Rfam: an RNA family database. Nucleic Acids Res. 31, 439–41 (2003).

80. Nawrocki, E. P. & Eddy, S. R. Infernal 1.1: 100-fold faster RNA homology searches. Bioinformatics 29, 2933–5 (2013).

81. Löytynoja, A. Phylogeny-aware alignment with PRANK. in Methods in molecular biology (Clifton, N.J.) 1079, 155–170 (2014).

82. Nguyen, L. T., Schmidt, H. A., Von Haeseler, A. & Minh, B. Q. IQ-TREE: A fast and effective stochastic algorithm for estimating maximum-likelihood phylogenies. Mol. Biol. Evol. 32, 268–274 (2015).

83. Kalyaanamoorthy, S., Minh, B. Q., Wong, T. K. F., von Haeseler, A. & Jermiin, L. S. ModelFinder: fast model selection for accurate phylogenetic estimates. Nat. Methods 14, 587–589 (2017).

84. Hoang, D. T., Chernomor, O., von Haeseler, A., Minh, B. Q. & Vinh, L. S. UFBoot2: Improving the Ultrafast Bootstrap Approximation. Mol. Biol. Evol. 35, 518–522 (2018).

85. Han, M. V., Thomas, G. W. C., Lugo-Martinez, J. & Hahn, M. W. Estimating Gene Gain and Loss Rates in the Presence of Error in Genome Assembly and Annotation Using CAFE 3. Mol. Biol. Evol. 30, 1987–1997 (2013).

86. Mitchell, A. L. et al. InterPro in 2019: improving coverage, classification and access to protein sequence annotations. Nucleic Acids Res. (2018). doi:10.1093/nar/gky1100

87. Altschul, S. F., Gish, W., Miller, W., Myers, E. W. & Lipman, D. J. Basic local alignment search tool. J. Mol. Biol. 215, 403–410 (1990).

88. Krogh, A., Larsson, B., Von Heijne, G. & Sonnhammer, E. L. Predicting transmembrane protein topology with a hidden markov model: application to complete genomes11Edited by F. Cohen. J. Mol. Biol. 305, 567–580 (2001).

89. Katoh, K. & Standley, D. M. MAFFT Multiple Sequence Alignment Software Version 7: Improvements in Performance and Usability. Mol. Biol. Evol. 30, 772–780 (2013).

90. Capella-Gutiérrez, S., Silla-Martínez, J. M. & Gabaldón, T. trimAl: A tool for automated alignment trimming in large-scale phylogenetic analyses. Bioinformatics 25, 1972–1973 (2009).

91. Suyama, M., Torrents, D. & Bork, P. PAL2NAL: Robust conversion of protein sequence alignments into the corresponding codon alignments. Nucleic Acids Res. 34, (2006).

92. Smith, M. D. et al. Less is more: an adaptive branch-site random effects model for efficient detection of episodic diversifying selection. Mol. Biol. Evol. 32, 1342–53 (2015).

93. Pond, S. L. K., Frost, S. D. W. & Muse, S. V. HyPhy: hypothesis testing using phylogenies. Bioinformatics 21, 676–679 (2005).

